# Different armpits under my new nose: olfactory sex but not gender affects implicit measures of embodiment

**DOI:** 10.1101/2021.12.16.472829

**Authors:** Marte Roel Lesur, Yoann Stussi, Philippe Bertrand, Sylvain Delplanque, Bigna Lenggenhager

**Affiliations:** Department of Psychology, University of Zurich, Zurich, Switzerland; Department of Psychology, Harvard University, Cambridge, MA, USA; Swiss Center for Affective Sciences, University of Geneva-CISA, Geneva, Switzerland; Institute of Psychology, Faculty of Humanities and Social Sciences, Université de Paris; BeAnotherLab, Spain

**Keywords:** Olfaction, Embodiment, Gender, Plasticity, Multisensory Integration, Bodily Illusion

## Abstract

Research has shown that conflicting multisensory signals may alter embodiment to the point of self-identifying with a foreign body, but the role of olfaction in this process has been overlooked. Here, we study in healthy participants how sex (male and female sweat odors) and gender (male and female cosmetic scents) olfactory stimuli contribute to embodiment. Participants saw from the perspective of a sex mismatching person in virtual reality and received synchronous visuo-tactile stimulation to elicit illusory embodiment of the seen body while smelling either sex- or gender-congruent stimuli. We assessed implicit (skin conductance responses to visual threats) and explicit (questionnaire) measures of embodiment. Stronger responses to threat were found when participants smelled the sex-congruent compared to the sex-incongruent odor, while no such differences were found for the cosmetic scents. According to the questionnaire, embodiment did not differ between conditions. Post-experimental assessment of the presented cues, suggest that while both sweat odors were considered generally male, cosmetic scents were not. The presented scents were generally not associated to the embodied body. Our results suggest that sex-related body odors influence implicit but not explicit aspects of embodiment and are in line with unique characteristics of olfaction in other aspects of cognition.

## Introduction

‘Who smelt it, dealt it’ is a phrase used in several languages to indicate ownership and agency of a flatulence. Despite similar linguistic examples that convey messages about the link of olfaction and ownership, the self, or interpersonal relations (e.g., ‘I cannot smell this person’ when disliking someone in several languages), we often seem to disregard the role of olfaction when scientifically studying human self-identity. This notably contrasts with experimental work in nonhuman animals; for instance, dogs, while not passing the well-known mirror-self recognition task (Gallup, 1970), have been shown to successfully recognize their own scent in an olfactory equivalent of the task (Horowitz, 2017). As is the case with canines, odors are an integral part of animal interactions and guide individual behavior across a wide variety of species. In animal research, it has been long accepted that odors mediate basic behaviors such as kin recognition, sexual identification, and sexual attraction (Russell, 1976). By comparison, the fact that humans are able to extract biological and social cues from body odors has long been dismissed outright (Lundström & Olsson, 2010). Yet, humans permanently produce and process body odors, even if largely on a subconscious level (Perl et al., 2020; Prehn et al., 2006; Zhou & Chen, 2008, 2009). These odors seem to play a role in the perception of self and others and in self-other distinction (Perl et al., 2020). Humans can indeed distinguish their own body odor from that of other persons’. Body as compared to non-body odors in humans are linked to specific neural substrates including multisensory integration areas such as the angular gyrus (Lundström et al., 2008), a network associated to bodily self-perception (Blanke et al., 2002, 2004). One’s own body odor is considered to be a quite stable and unique “signature” of the bodily self (Lundström & Olsson, 2010), and it has been argued that humans partake in self-smelling behaviors as a way of reassuring the self (Perl et al., 2020). In fact, body odors deliver a great range of information about the person, from individual/kin identity, age, illness, reproductive state, attraction, to transient emotional states (Chen & Haviland-Jones, 2000; Ferdenzi et al., 2020; Mallet & Schaal, 1998; Porter, 1998, 1998; Semin & Groot, 2013, for a review).

On this basis, it seems reasonable to assume that the perception of one’s own body odor is integrated with other senses (e.g., visual, tactile, motor, auditory, interoceptive, nociceptive, vestibular) to create and maintain a coherent sense of a bodily self. Bodily self-consciousness is thought to be dependent on a continuous integration and updating of signals from various modalities and prior beliefs (Apps & Tsakiris, 2014; Blanke et al., 2015). It has been experimentally studied by providing participants with spatiotemporally congruent multisensory signals, which are mediated by top-down associations with the embodied object. Stimulation to both their hidden own and a seen virtual or fake full-bodies or body parts results in illusory embodiment and self-identification with the foreign body or limb despite clear morphological differences from the own body (Botvinick & Cohen, 1998; Michel et al., 2014; Petkova & Ehrsson, 2008; Slater et al., 2010; see Kilteni et al., 2015 for a review), pointing at the remarkable malleability of the sense of a bodily self. In fact, experimental alterations of the bodily self have been shown to affect physiological, behavioral, affective, and cognitive processes (e.g., Banakou et al., 2018; Falconer et al., 2014; Maister et al., 2015; Moseley et al., 2008; Peck et al., 2013). However, while many studies used multisensory conflicts relying on a/synchrony between signals coming from different combinations of sensory modalities (Aspell et al., 2013; Capelari et al., 2009; Ehrsson et al., 2005; Kannape et al., 2010; Macauda et al., 2014; Tajadura-Jiménez et al., 2012; Tsakiris et al., 2006; Walsh et al., 2011), the contribution of own-body olfactory cues to illusory embodiment and self-identification has, to our best of knowledge, not yet been addressed. Recent studies have shown that (a) visuo-olfactory congruent cues result in a higher degree of self-identification with an arbitrary object (a grapefruit) seen in the position of the own body (Roel Lesur, Aicher, et al., 2020), and (b) feeling of one’s own body was reported as lighter or heavier depending on the concurrent scent (Brianza et al., 2019), thereby suggesting that prior associations of odors might influence various aspects of the sense of bodily self.

Here we investigated the effects of sex- (artificial female/male sweat odor) and gender- (female/male stereotypical cosmetic odor) congruent/incongruent body-related odors on illusory embodiment in virtual reality. Differences between sexes in the composition of sweat are supported by distinct chromatographic profiles of volatile compounds (Penn et al., 2007), non-volatile odor-precursors (Troccaz et al., 2009), as well as varying amounts of axillary skin microflora (Jackman & Noble, 1983). This evidence is in line with research suggesting that bodily chemical compounds signal sex-specific cues (Gustavson et al., 1987; Lundström et al., 2006; Olsson et al., 2006; Wyart et al., 2007). Such compounds have been shown to modulate cognitive processes such as gait perception depending on the participants’ sex and sexual preference (Ye et al., 2019) or masculinity and femininity ratings of body movements (Zhou et al., 2014). While sex-specific compositions of odors are at least partially biologically-driven given their dependency on genetic and anatomical factors (Jackman & Noble, 1983; Savelev et al., 2008), cultural associations can also determine certain scents in a gender-specific manner, as has been stereotyped by cosmetics (Donna, 2009; Lindqvist, 2013).

In the current study, participants were blindly exposed to either sex- (female/male sweat odor; corresponding to body odors) or gender- (female/male stereotypical cosmetic odor; corresponding to cosmetic scents) congruent or incongruent odors in an embodied virtual reality setup. While exposed to the scents, they saw from the perspective of a previously recorded sex-mismatched person in a head mounted display (HMD) and experienced congruent visuo-proprioceptive and visuo-tactile stimulation, which have been consistently used to induce embodiment illusions (Kilteni et al., 2015) including embodiment of different genders (Bertrand et al., 2014; Bolt et al., 2021; Neyret et al., 2020; Petkova & Ehrsson, 2008; Seinfeld et al., 2018; Tacikowski et al., 2020). At different moments of the stimulation procedure, a visual threat was presented to the seen body, and electrodermal response to threat was measured for each of these events (Armel & Ramachandran, 2003; Petkova & Ehrsson, 2008; Preuss & Ehrsson, 2019). Explicit illusory embodiment was assessed through a questionnaire after each condition. We expected stronger self-identification as measured by explicit and implicit measures in the visuo-olfactory congruent as compared to the incongruent blocks. Further explorative measures were taken to assess the participants’ perception of the presented odors in a second experimental block.

## Methods

### Participants

Based on sample sizes from similar studies on virtual-reality based bodily illusions (e.g., Kokkinara et al., 2016; Lenggenhager et al., 2007; Maselli & Slater, 2013; Slater et al., 2009), 24 participants without any history of psychiatric or neurological disorders were recruited at the University of Zurich and received either university credits or a financial compensation (20 CHF) for their participation. They provided written informed consent. The experimental protocol was approved by the Ethics Committee of the Faculty of Arts and Social Sciences at the University of Zurich (Approval Number 17.12.15) and followed the ethical standards of the Declaration of Helsinki. Two participants interrupted the procedure and thus their data was removed from subsequent analyses, with the final sample consisting of 22 participants (12 females) ranging between 18 and 42 years old (*M* = 28.8, *SD* = 7.5). Electrodermal activity data from two additional participants was discarded due to data loss, resulting in a sample size of 20 participants for this measure (N = 20; 11 females; age *M* = 29, *SD* =7.6).

### Apparatus

#### Visuotactile stimulation

An Oculus CV1 HMD was used for stimulation. The software was designed using Unity 2018.2.8 for displaying a 235-degree prerecorded video portraying the first-person perspective of a real male or female sex person. The software ran on a computer (Nvidia Geforce GTX 1080 8 GB; 16 GB RAM; Intel Core i7, 3.2 GHz) connected to the HMD. The videos were respectively filmed from the perspective of an actor and an actress, using a monoscopic Kodak SP360 4K camera at a resolution of 2,160 × 2,160 pixels at 30 frames per second. Another sex-matched person was recorded from a third person perspective interacting with them, this person synchronized their movements and actions to a previously recorded set of audio instructions to achieve consistent timing between videos. The same audio was later heard by the experimenter during the stimulation procedure to imitate those movements in synchrony with the video (see Figure 1). Additional visual monitoring on a computer further facilitated this synchronization for the experimenter. This protocol is based on a method developed by BeAnotherLab and widely used in diverse settings (BeAnotherLab, 2020; Bertrand et al., 2014; Sutherland, 2015) including several scientific studies (Bertrand, 2021; Roel Lesur, Lyn, et al., 2020; Roel Lesur, Aicher, et al., 2020). The questionnaires were displayed on the HMD and answered by looking at a fixed position on the HMD with a pointer for a period of 1 s.

**Figure 1.**
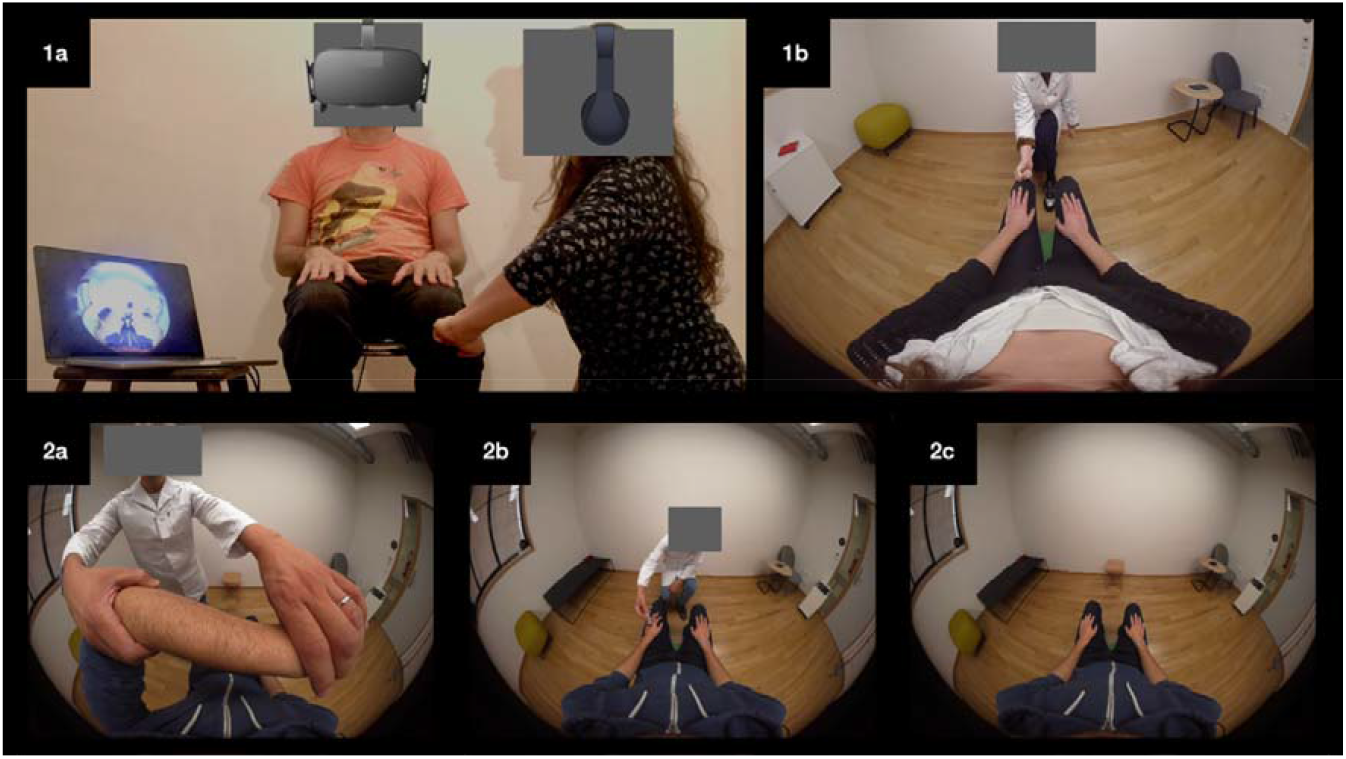
**1a)** illustrates a male participant wearing the HMD with an olfactory pen attached to it while receiving tactile stimulation from the experimenter. The experimenter is wearing headphones where the timing for the tactile stimulation is presented while at the same time visually monitoring the gestures. This image was recreated for illustration purposes. **1b)** depicts what is seen at this moment by the male participant on the HMD. **2a, 2b and 2c** depict frame captures from the immersive videos presented on the HMD to female participants. The first illustrates the initial gesture of their own arm being brought towards their nose, the second shows a threat (a syringe pinching the hand), and the third depicts the static sitting position. For the purposes of this preprint, the faces have been covered to hide any features that may identify the people involved,.

**Figure 2.**
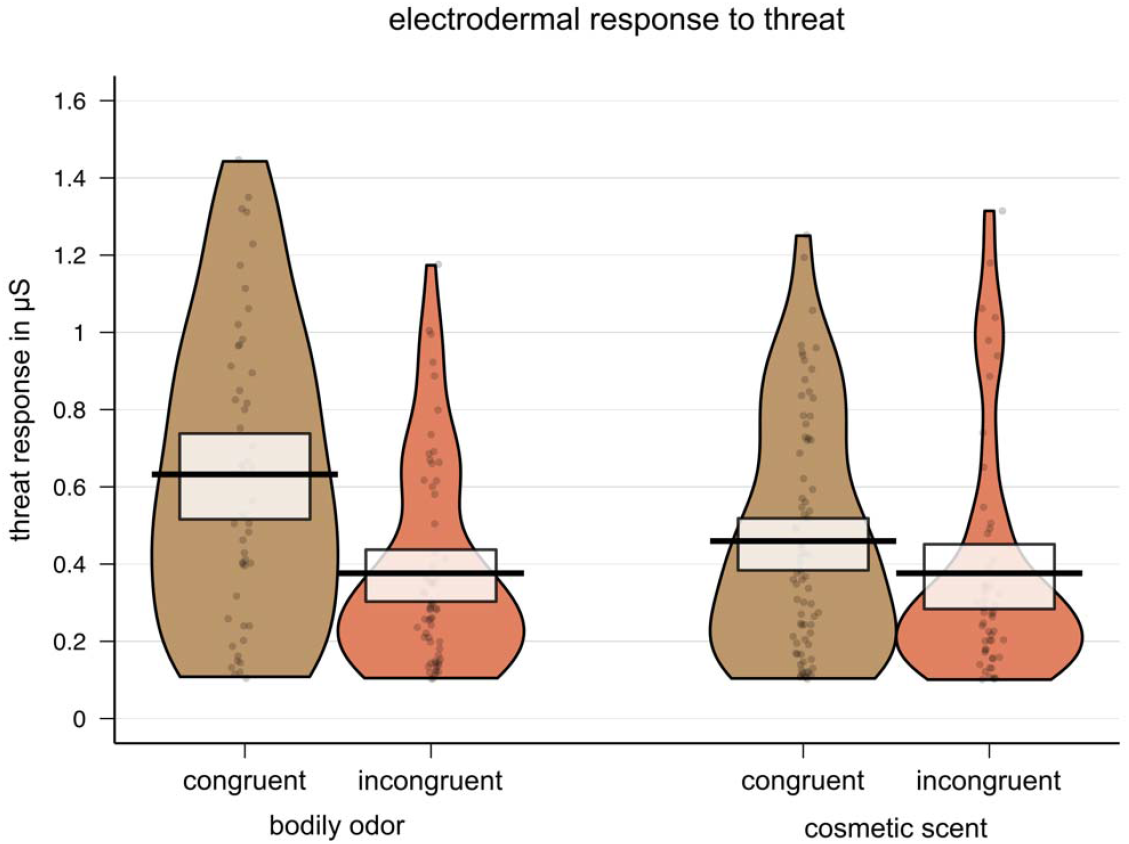
Raw values, central tendencies and distribution of skin conductance responses grouped by congruency and odor type.

#### Olfactory stimulation

Three different compounds [MSH (3-methyl-3-sulfanylhexan-1-ol), HMHA (3-hydroxy-3-methylhexanoic acid) and M2HA (3-methyl-2-hexenoic acid)] were used to create two synthesized sweat mixtures that respect sexual dimorphism observed for real sweat (Troccaz et al., 2009). Female synthesized sweat was composed of 80% HMHA, 10% M2HA and 10% MSH and diluted 1000 times in triacetin (female sweat odor). Male synthesized sweat was composed of 80% HMHA and 20% M2HA and diluted 100 times in triacetin (male sweat odor). A typically feminine scent (Chloé eau de Parfum at 10% in dipropylene glycol; DIPG; female cosmetic scent), and masculine pleasant scent (Hugo Deep Red at 10% in DIPG; male cosmetic scent) were used as pleasant body-related scents. Two extra scents were used for the odor perception block (mushroom 10% in DIPG and rose 10% in DIPG, respectively), these were previously selected as a rather negative scent and rather positive scent based on tests from a small-scale sample (see Supplementary Materials). All compounds were provided by Firmenich, S.A. The solutions (3 ml) were injected into the tampon of cylindrical felt-tip pens (14-cm long, inner diameter 1.3 cm). The use of these highly practical devices (provided by Burghart, Germany) avoids any contamination of the environment.

#### Skin conductance recordings

Threat-evoked skin conductance responses (SCRs) were measured using two Ag-AgCl electrodes (6 mm diameter contact area with a 1.6 mm cavity for electrode gel) mounted in individual polyutherane housings attached to the palmar side of the middle phalanges of the first and second digits of the participants’ right hand. Skin conductance was continuously recorded with a Biopac MP150 system (Santa Barbara, CA, USA) and EDA100C amplifier (Biopac Systems Inc., Goleta, CA, USA) at a 500 Hz sampling rate running in an additional Windows computer.

### Procedure

#### Embodiment

Upon arrival, participants were informed that they would see from the perspective of another body in virtual reality and that they would be touched on the arms, feet, and knees. They were told that a virtual threat would be presented but that there would be no real threat to their bodies. There was no reference to the odors at this stage. After providing written informed consent, the electrodes were placed, after which participants were helped to put on the HMD and sat down. They were required to put their hands on their laps and their feet on the ground to match their posture to that seen on VR (see Figure 1). A single test question was presented to confirm that participants understood how to answer the questionnaire on a visual analogue scale (VAS) by facing towards the desired location on the HMD. Before beginning the stimulation, for each condition, a pen with synthesized odor was fixed on to the HMD just below the nose (see Figure 1). This pen contained one of four synthesized odors, namely male or female cosmetic scent, or male or female sweat odor that were presented in a counterbalanced manner. Participants then saw a sex-mismatching person from a first person-perspective on the video. The videos started with an experimenter grabbing the left arm seen from the participants’ perspective and taking the elbow close to the nose (this was intended as a simile to smelling the armpit). Other than this passive movement directed by the experimenter (Figure 1, 2a), participants were sitting still with their hands on their laps (Figure 1, 2c). A series of eight visuotactile stimuli (to enhance embodiment) and seven visual threats (to measure implicit embodiment; see Figure 1, 2b) were presented during each video. For both male and female participants, there were two videos with differing orders for the threatening stimuli and visuotactile stimulation. The order of these was semi-counterbalanced to minimize potential order effects. Participants saw each sex-mismatched video twice, once per condition, and each condition was followed by an embodiment questionnaire.

#### Odor perception

After finalizing the above procedure, participants were presented with each of the four scents applied in the previous block plus two extra scents (mushroom and rose, respectively) in a counterbalanced manner. The additional two scents were included as control items, and one was as rather negative (mushroom), while the other as rather positive (rose) in a small naive sample (see Supplementary Materials). In the main study, participants were instructed to smell each of the six scents for 10 s, while they were blindfolded. A questionnaire consisting of interleaved forced choice and VAS responses followed to assess the recognition, pleasantness, and gender-association of each scent. A debriefing questionnaire consisting of 5 questions was presented on the HMD upon completion of all the conditions. These questionnaires were responded using a computer and a mouse.

### Measures

#### Explicit embodiment measure: questionnaire

The embodiment questionnaire was adapted from previous studies (Botvinick & Cohen, 1998; Dobricki & Rosa, 2013; Gonzalez-Franco & Peck, 2018; Lenggenhager et al., 2007) and answered on a VAS ranging from *strongly disagree* (0) to *strongly agree* (1) on the HMD through head movements. The embodiment questionnaire items are presented in Table 1.

**Table 1.** Results of ART ANOVAs for the embodiment questionnaire (N =22).

| Item | Congruency |  |  |  |  | Odor Type |  |  |  |  | Interaction |  |  |  |  |
| --- | --- | --- | --- | --- | --- | --- | --- | --- | --- | --- | --- | --- | --- | --- | --- |
| | *F*(df) | *p* | $\eta^2$ | 90% CI | | *F*(df) | *p* | $\eta^2$ | 90% CI | | *F*(df) | *p* | $\eta^2$ | 90% CI | |
| I felt comfortable in the room | 0.33 (1,21) | .57 | .02 | .00 | .19 | 0.01 (1, 21) | .94 | .00 | .00 | .01 | 0.13 (1, 21) | .73 | .01 | .00 | .15 |
| Sometimes the body I saw when looking down felt like a stranger | 2.6 (1,21) | .12 | .11 | .00 | .34 | 0 (1, 21) | .98 | .00 | .00 | .00 | 2.64 (1, 21) | .12 | .11 | .00 | .34 |
| Sometimes it felt like I could control the seen body like my own. | 0.23 (1,21) | .63 | .01 | .00 | .17 | 0.45 (1, 21) | .51 | .02 | .00 | .20 | 0.28 (1, 21) | .60 | .01 | .00 | .18 |
| Sometimes it felt as if the touches I felt were caused by the ones I saw | 0.7 (1,21) | .41 | .03 | .00 | .23 | 0.11 (1, 21) | .74 | .01 | .00 | .14 | 0.17 (1, 21) | .68 | .01 | .00 | .16 |
| Sometimes it felt as if the body I saw was like my own in terms of shape or skin color or appearance | 1.51 (1,21) | .23 | .07 | .00 | .28 | 2 (1, 21) | .17 | .09 | .00 | .31 | 0.06 (1, 21) | .81 | .00 | .00 | .11 |
| Sometimes it felt like I was in the middle of the action. | 0.39 (1,21) | .54 | .02 | .00 | .19 | 0.31 (1, 21) | .58 | .02 | .00 | .18 | 0.59 (1, 21) | .45 | .03 | .00 | .22 |
| Sometimes I had the feeling that I could be threatened by what I saw | 0.23 (1,21) | .64 | .01 | .00 | .17 | 0.09 (1, 21) | .77 | .00 | .00 | .13 | 0.1 (1, 21) | .75 | .01 | .00 | .01 |
| Sometimes it felt like I had a male/female body | 0.6 (1,21) | .45 | .03 | .00 | .22 | 0.88 (1, 21) | .36 | .04 | .00 | .24 | 0.8 (1, 21) | .38 | .04 | .00 | .24 |
| Sometimes I felt as if I could move the seen body | 0 (1,21) | .96 | .00 | .00 | .00 | 1.12 (1, 21) | .30 | .05 | .00 | .26 | 0.03 (1, 21) | .87 | .00 | .00 | .09 |
| I felt good in my body | 1.73 (1,21) | .20 | .08 | .00 | .30 | 0.93 (1, 21) | .35 | .04 | .00 | .25 | 0.01 (1, 21) | .93 | .00 | .00 | .03 |
| Sometimes it felt like I was wearing different clothes than before the experiment | 1.91 (1,21) | .18 | .08 | .00 | .31 | 0.31 (1, 21) | .58 | .02 | .00 | .18 | 1.18 (1, 21) | .29 | .05 | .00 | .26 |
| Sometimes the body I saw felt like my own | 2.78 (1,21) | .11 | .12 | .00 | .35 | 0.04 (1, 21) | .85 | .00 | .00 | .00 | 1.4 (1, 21) | .25 | .06 | .00 | .28 |
| In general, the touches I saw and felt were synchronous | 0.05 (1,21) | .82 | .00 | .00 | .11 | 1.69 (1, 21) | .21 | .07 | .00 | .29 | 1.31 (1, 21) | .27 | .06 | .00 | .27 |

#### Implicit embodiment measure: skin conductance response to threat

For synchronizing the virtual reality stimulation with the physiological recordings, triggers were sent through a serial port from the stimulation computer to the Biopac system using a trigger box. The triggers were previously timed to correspond to the first moment at which the visual threat appears on the screen. A total of 7 threatening stimuli were included per condition, each corresponding to a direct bodily threat to the seen body, respectively as follows: a knife stabbing the leg, a syringe injecting the arm, a hammer hitting the foot, and a ball being thrown at the belly. The first three of these items were repeated two times, once on the left and once on the right limb in such a way that two subsequent threats would not be on the same side or limb. The ball thrown at the belly was presented only once.

Specific SCRs were measured in microSiemens (μS) and analyzed offline using Ledalab software (v3.4.9, Benedek & Kaernbach, 2010). For each trial, SCRs were scored as the peak-to-peak amplitude difference in skin conductance of the response falling within −2 to 5 s of the trigger onset. The minimal response criterion was set as 0.01 μS and only trials where there was a single response in the temporal window of interest were included. The raw SCR scores were square-root-transformed to normalize the distributions. Because the inclusion of zero responses (i.e., using SCR magnitude) is subject to confounds between the response strength and the frequency (Dawson et al., 2016; Preuss & Ehrsson, 2019; Prokasy & Raskin, 1973) and the large number of stimuli and repetitions in our setup, we here excluded zero responses from analysis (i.e., using SCR amplitude). This was motivated by the fact that we were specifically interested in assessing threat-related responses, which are a standard measure in the investigation of bodily illusions (Armel & Ramachandran, 2003; Ehrsson, 2007; Petkova & Ehrsson, 2008; Preuss & Ehrsson, 2019) but are well known to be reduced after several repetitions (Dawson et al., 2016; Preuss & Ehrsson, 2019). Such measure of SCR amplitude has been used as an indicator of illusion strength in related research (e.g., Preuss & Ehrsson, 2019).

#### Odor perception

A questionnaire with interleaved forced choice and VAS items was included after participants smelled each stimulus. The questionnaire was responded on a computer screen using a mouse and included the items presented on Table S1 (Supplementary Materials).

#### Debriefing

A final debriefing questionnaire was presented on a computer screen and answered on a VAS ranging from *strongly disagree* (0) to *strongly agree* (1). The items are listed in the following Results section.

### Statistical analyses

Statistical analyses were conducted in R version 3.5.1 (R Core Team, 2020) and inspected for normality through both visual inspection and the corresponding statistical tests. Two-tailed comparisons are reported.

Data from the embodiment and odor perception questionnaires were analyzed with separate aligned rank transformation (ART) ANOVAs using the package ARTool (Kay & Wobbrock, 2020) for each item. We used ART ANOVAs rather than conventional ANOVAs because of the nonparametric nature of the questionnaire data and the robustness of ART ANOVAs for nonparametric analyses (Wobbrock et al., 2011). Statistically significant findings were followed up with multiple Bonferroni-corrected Wilcoxon signed-rank for the comparisons of interest. We report median and interquartile range (IQR) as descriptive statistics for nonparametric data, as well as partial eta-squared (η_p_^2^) as an estimate of effect size together with the corresponding 90% confidence intervals (CI).

SCR data was analyzed by means of a general linear mixed-effects model using the lme4 (Bates et al., 2015) and lmerTest (Kuznetsova et al., 2017) packages. We entered the within-participants factors congruency (congruent vs. incongruent) and odor type (sweat odor vs. cosmetic scent) and their interaction as fixed effects. As random effects, we modeled random intercepts for participants and by-participant random slopes for congruency and odor type. The by-participant random slope for the interaction was not included in the random-effects structure, as its inclusion led to model singularity, indicating overfitting (Bates et al., 2018). A principal component analysis of the random-effects covariance matrix estimates (Bates et al., 2018) confirmed that the inclusion of the by-participant random slope for the interaction led to overfitting in returning four principal components, whereas three were sufficient to account for 100% of the variance explained (i.e., the fourth component explained 0% of the random-effects variance). The final model was built as follows (in lme4 syntax): *sqrtSCR ~ (congruency * odor.type) + (1 + congruency + odor.type|participant)*. We used the ‘bobyqa’ optimizer and set the number of model iterations to 200’000 to fit the model. Follow-up comparisons were computed with the emmeans (Lenth, 2020) package when appropriate and Bonferroni correction was applied to correct for multiple testing. We report Cohen’s *d* for linear mixed-effects models (see Brysbaert & Stevens, 2018; Westfall et al., 2014) as an estimate of effect size and their 95% CI.

## Results

### Embodiment questionnaire

The ART ANOVA performed for each item of the revealed no statistically significant differences between conditions for any items of the questionnaire (see Table 1).

### Skin conductance response to threat

A Bayesian analysis of variance (ANOVA) using JASP version 0.11.1 (JASP Team) showed moderate evidence for a similar repartition of the zero responses across the various conditions of the congruency (BF_01_ = 3.88) and the odor type (BF_01_ = 3.52) factors, along with their interaction (BF_01_ = 14.15). This suggests that there were no differences in the proportion of zero responses across the experimental conditions.

Results from the linear mixed-effects model conducted on the SCR data are reported in Table S3 (Supplementary Materials). In line with our prediction that visuo-olfactory sex congruence modulates threat-evoked SCRs as an implicit measure of embodiment, we found a statistically significant main effect of congruency, indicating that participants showed higher SCR amplitude to threat when they were exposed to a sex- or gender-congruent odor (*M* = 0.52, *SE* = 0.05) than to a sex- or gender-incongruent odor (*M* = 0.36, *SE* = 0.05), *b* = .08, 95% CI [.005, .15], *p* = .028, *d* = 0.512, 95% CI [0.036, 0.988]. The main effect of odor type conversely did not yield statistical significance, *b* = .02, 95% CI [-.04, .08], *p* = .387, *d* = 0.161, 95% CI [-0.252, 0.574].

Importantly, the main effect of congruency was qualified by the higher-order interaction between congruency and odor type, *b* = .05, 95% CI [.01, .09], *p* = .005, *d* = 0.700, 95% CI [0.181, 1.220]. Follow-up comparisons revealed that participants exhibited higher SCR amplitude to threat when exposed to a sex-congruent (*M* = 0.60, *SE* = 0.08) than a sex-incongruent (*M* = 0.33, *SE* = 0.05) sweat odor, *t*(34.8) = 3.23, *p* = .005, *d* = 0.862, 95% CI [0.315, 1.410]. By contrast, no such difference emerged when participants were exposed to a gender-congruent (*M* = 0.44, *SE* = 0.05) versus gender-incongruent cosmetic scent (*M* = 0.39, *SE* = 0.06), *t*(29.5) = 0.64, *p* > .99, *d* = 0.162, 95% CI [-0.358, 0.683]. This suggests that participants’ implicit embodiment, as measured with threat-evoked SCR amplitude, was specifically enhanced when they were exposed to a sex-congruent sweat odor compared to a sex-incongruent sweat odor, but not when they were exposed to gender-congruent and incongruent cosmetic scents.

### Odor perception questionnaire

For each of the VAS items of the odor perception questionnaire (see Table S1), an ART ANOVA was performed including the single factor condition with six levels, each corresponding to each of the presented odors. Statistically significant differences were found for the *gender attribution* (*F*(5, 105) = 9.06, *p* < .001, η_p_^2^ = .302, 90% CI [0.16, 0.39]), *liking* (*F*(5, 105) = 18.46, *p* < .001, η_p_^2^ = .468, 90% CI [0.34, 0.55]) and *intensity* (*F*(5, 105) = 3.07, *p* = .013, η_p_^2^ = .127, 90% CI [0.02, 0.2]) ratings of the odors, but not for the *familiarity* ratings (*F*(5, 105) = 0.41, *p* = .84, η_p_^2^ = .019, 90% CI[0.00, 0.029]). Bonferroni-corrected multiple comparisons between the relevant odors for the significant items are reported in Table 2. The forced-choice items were analyzed using binomial tests to assess whether the proportion of *yes/no* responses was different from chance (50%). Results from these analyses are presented in Table 3.

**Table 2.**
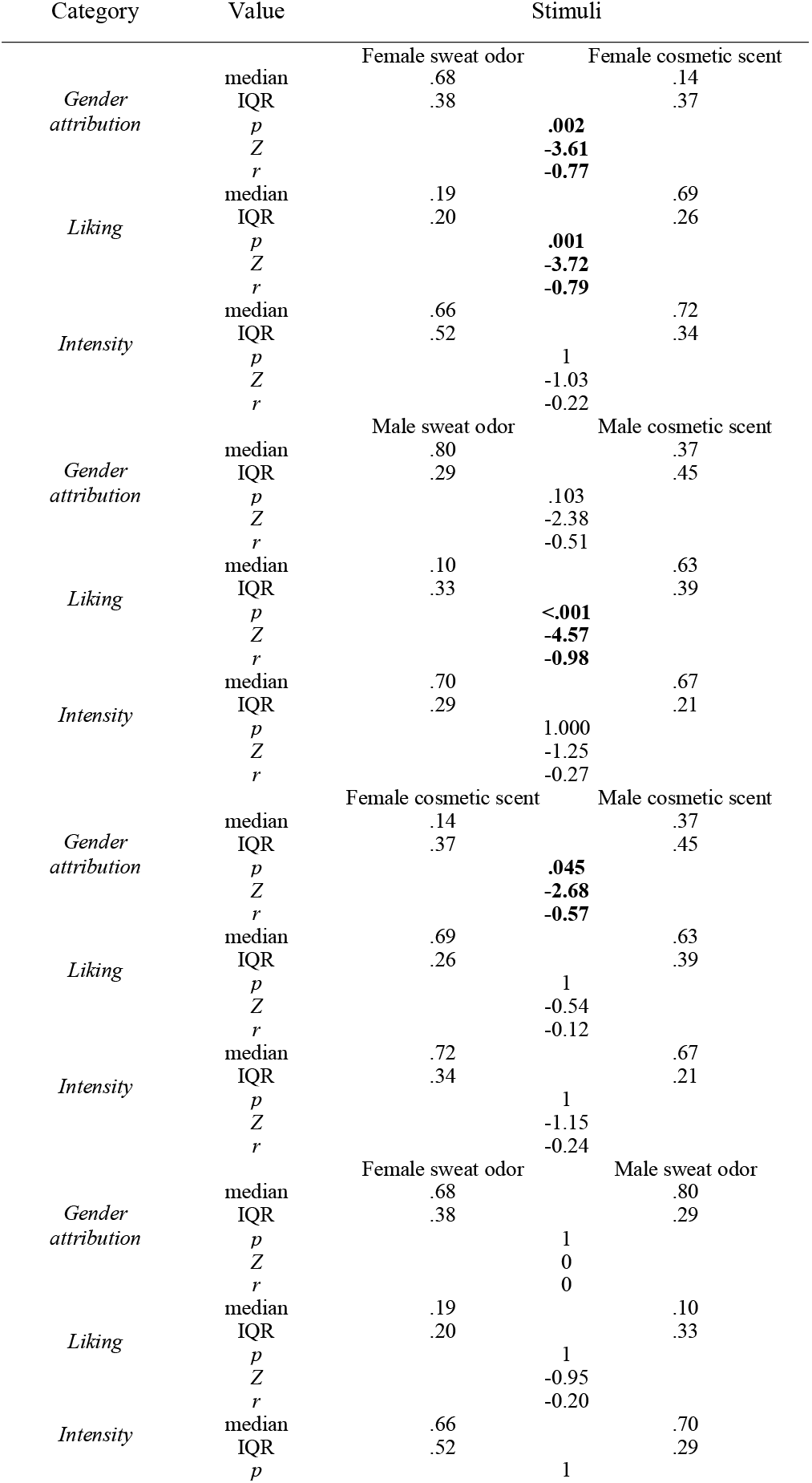
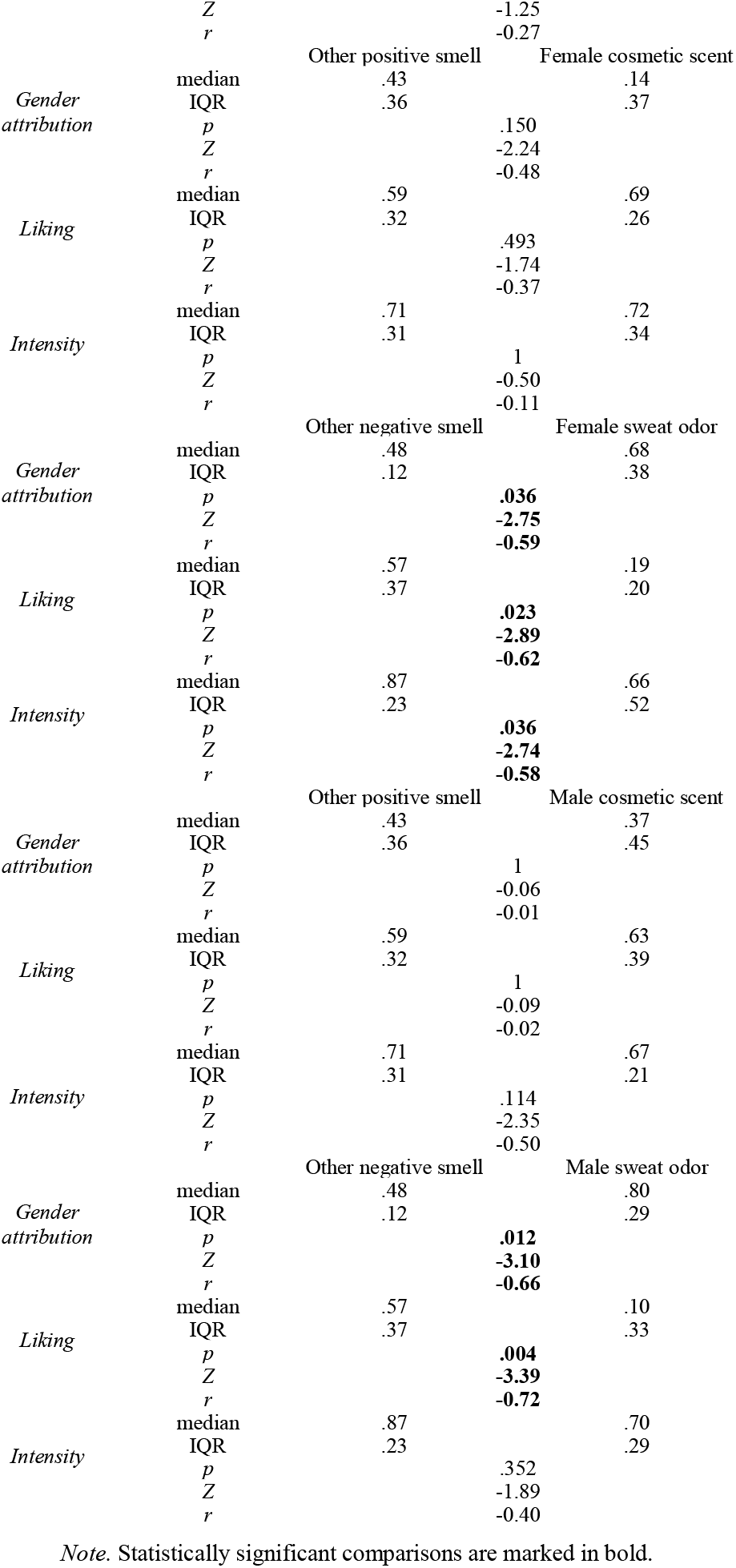
Bonferroni-corrected multiple comparisons for the VAS items of the odor perception questionnaire as well as median and IQR for each stimulus and category

**Table 3.**
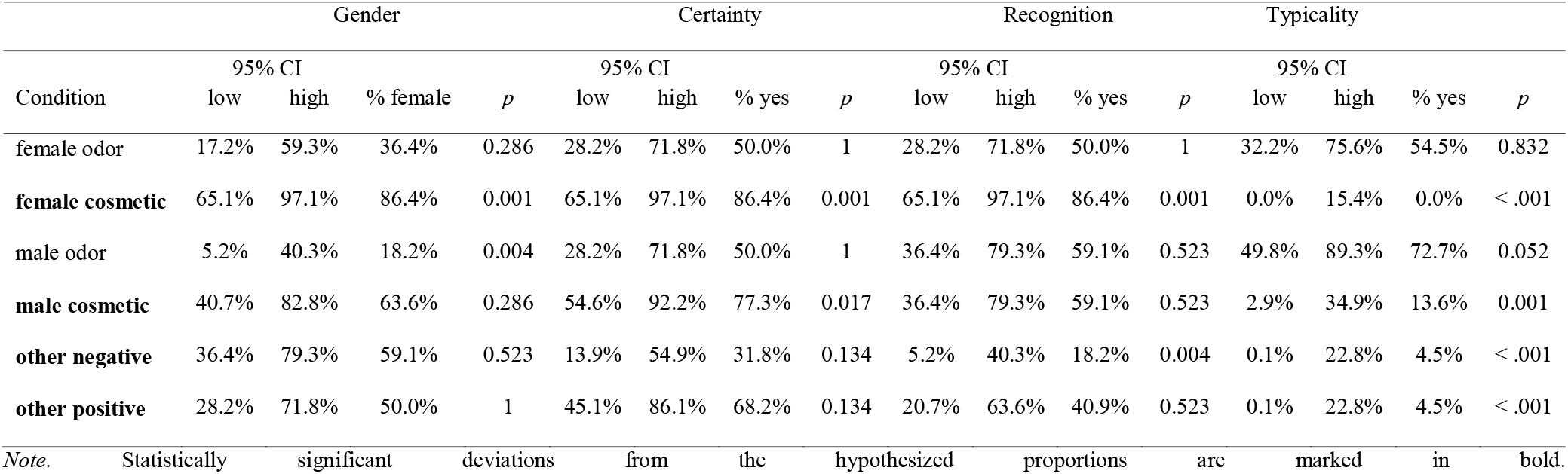
Multiple binomial tests for the forced choice items of the odor perception questionnaire with a hypothesized proportion of 50%.

### Debriefing

Each of the debriefing questions yielded the following scores: *1) I thought that the odor was coming from the body that I saw in my position* (Median = 0.19, IQR = 0.35*); 2) During the VR film I paid attention to the smells* (Median = 0.41, IQR = 0.55); *3) The smell changed the way I felt about the body that I saw in my position* (Median = 0.21, IQR = 0.51); *4) I thought the smell was coming from the experimenter or the room I saw in the video* (Median = 0.54, IQR = 0.41); *5) The smell changed my mood* (Median = 0.36, IQR = 0.38). For items *1)* and *4)* we additionally calculated these scores excluding three participants that reported not recognizing any of the presented smells (Median = 0.24, IQR = 0.4 and Median = 0.51, IQR = 0.43, respectively).

## Discussion

Our study used a well-established virtual-reality-based protocol for eliciting illusory embodiment of a mismatching-sex body using a first-person perspective and synchronous visuo-tactile stimulation (Bertrand, 2021; Bertrand et al., 2014; Bolt et al., 2021; Petkova & Ehrsson, 2008; Tacikowski et al., 2020). As participants saw from the perspective of a sex-mismatching body, they were presented with gender- (cosmetic scents) and sex-related (sweat odors) olfactory stimuli that were either congruent or incongruent with the seen body. We aimed to evaluate the contribution of body smells to the embodiment of a seen body. Explicit illusory embodiment was assessed through a questionnaire, and implicit illusory embodiment through SCR. The former showed no differences between odor conditions; however, our implicit measure yielded stronger reactions for the congruent sweat odors (when the seen and smelled avatar were sex-congruent), compared to both cosmetic scents and incongruent sweat odors. A subsequent questionnaire on odor perception showed some important findings which might shed light on the underlying processes. First, while there were differences in gender-associations for the cosmetic scents (with the female scent being judged as more feminine), there were no differences between the sweat odors (with both being judged as rather male). Second, though a large majority of participants reported recognizing the female cosmetic scent from the stimulation procedure, this was not the case for the other olfactory stimuli. Lastly, most participants did not attribute the smell as emanating from the seen body. Together, these findings suggest that body odors might contribute to illusory embodiment at an implicit but not explicit level, which is in line with evidence suggesting unique patterns of olfaction (involving conscious content, post-perceptual processing, and memory, between other aspects) compared to the other major senses (e.g., Arshamian et al., 2020; Köster, 2002; Stevenson & Attuquayefio, 2013).

### Bodily self-identification and odors

Despite the relative lack of research directly addressing the role of smell in human bodily self-consciousness, it has been argued that we engage in subconscious self- or other-smelling behaviors as a process of reassuring the self (Perl et al., 2020). The behavior of self-smelling could thus be considered an analog of self-touch, which has been reckoned elemental in the development and healthy maintenance of our bodily self-consciousness (Husserl, 1952; Rochat, 1998; Roel Lesur et al., 2021). As Perl and colleagues (2020; *p*. 9) argue: “in sniffing our own body, we are subconsciously obtaining an external reflection and reassurance of *self*”. In our study, while participants did not explicitly state feeling a stronger sense of embodiment in the sex-congruent condition, their physiological reactions showed otherwise. When participants smelled the sex (sweat odor) but not gender (cosmetic scents) congruent stimuli, they showed stronger physiological reactions to threat. SCR is a common measure in the study of alterations of embodiment and bodily self-consciousness, where arguably our physiological responses are extended to a fake or virtual limb or full body when we self-identify with it (e.g. Armel & Ramachandran, 2003; Petkova & Ehrsson, 2008; Preuss & Ehrsson, 2019). This process is considered involuntary and subconscious (Armel & Ramachandran, 2003), and as such it is a relevant measure in practically all areas of psychology (Dawson et al., 2016). Our combined findings suggest no evident differences in explicit self-identification with the seen body, but differences in our implicit measure mediated by the concurrent odor. At this stage, we cannot conclusively attribute this modulation to subconscious processing (see e.g., Shanks et al., 2021), however further research might clarify the processes underlying our observations.

Previous research (Roel Lesur, Aicher, et al., 2020) reported explicit self-identification with an arbitrary object (a grapefruit) seen in virtual reality when there was a smell that was congruent with it (grapefruit scent) compared to an incongruent smell (strawberry scent). The reasons between explicit modulations of embodiment in the cited study and the lack of modulation here reported is unknown. It could be argued that in contrast to the study by Roel Lesur et al. (2020), the olfactory stimuli presented here were not as notable given that they are usual smells emanating from humans, compared to the fruity scents in the cited study. Alternatively, it could be that, in such study, the object seen in the location of the own body (i.e., a grapefruit) triggered no prior associations related to a human body, thus forcing participants to shift their attention to senses other than vision (i.e., smell) for grounding their sense of body. While in the cited study (Roel Lesur, Aicher, et al., 2020), the smells were identifiable (by a different experimental sample, as citric and strawberry, respectively), here the olfactory stimuli that yielded significant differences in SCRs (female and male sweat odors) were not judged as different in terms of the measured categories (gender, liking, recognition, and intensity). Furthermore, a previous study in a large sample showed that the bodily compounds used for this study were not explicitly distinguished by participants (Ferdenzi & Delplanque, 2021). Our integrated findings suggesting differences in explicit (odor-distinction and self-identification) and implicit measures might reflect a characteristic feature of olfaction that we discuss below.

### Unconscious processing of odors and implicit differences

Contrasting evidence between explicit and implicit measures has been often reported in the literature on bodily self-consciousness (e.g. de Haan et al., 2017; Roel Lesur et al., 2021; Roel Lesur, Weijs, et al., 2020; Rohde et al., 2011, 2013). Such divergent outcomes may seem surprising for stimulation through other modalities; nonetheless, in the context of olfaction, this might be in line with specific characteristics of the sense of smell. Olfactory processing in humans has been highlighted due to unique features in comparison with the other major senses. Amongst these, the conscious access to smells in terms of involuntary habituation, conscious content, attentional control, post-perceptual processing, and memory, seem to vary in contrast with other senses (Arshamian et al., 2020; Köster, 2002; Stevenson & Attuquayefio, 2013; Zucco, 2003). In fact, the subconscious processing of smells has been noted for its capacity to shift behavior (Gustavson et al., 1987; Holland et al., 2005; Mas et al., 2019; Olsson et al., 2006). Several studies have shown correct recollection behaviors without explicit recognition (Degel et al., 2001; Degel & Köster, 1999; Olsson & Cain, 2003) and findings suggest that odors alter cognition and behavior largely at a subconscious level (Prehn et al., 2006; Wisman & Shrira, 2015; Ye et al., 2019; Zhou et al., 2014; Zhou & Chen, 2008). This is, even when odors are not consciously perceived, they do impact our mood and behavior (Köster, 2002). Regarding self-other distinction, a study showed that participants were able to distinguish their own and their friends’ odors in a three-alternative forced choice task, but only with strikingly low confidence (Lundström et al., 2008). Another study showed no explicit recognition of one’s own body odor when compared to others in an experimental setting; however, disgust ratings were lower for the own odor (Übel et al., 2017). This supports the notion that despite a lack of clear direct identification, indirect psychological reactions support implicit olfactory self-other distinction. Following this, it does not seem implausible that despite the lack of changes in explicit self-identification, body odors did play a role in altering physiological responses related to embodiment, as our findings suggest.

A similar exclusive modulation of implicit embodiment measures has previously been found in other less attended senses like the vestibular system (Macauda et al., 2014). Our findings showed stronger SCRs for sex-congruent compared to sex-incongruent sweat odors, but this was not the case between gender-congruent and incongruent cosmetic scents. The reason behind this is not clear. However, this might suggest a potential implicit association to biologically-determined chemical compounds related to sex, which might be rooted in our cognitive system beyond cultural conceptions of gender. Alternatively, a hedonic bias triggering self-odor associations only for negatively perceived odors might explain these findings. One could imagine that when smelling sweat, a concern that this odor might emanate from oneself could trigger self-reassuring processes that are not activated for positive scents. At this point, these lines of reasoning remain purely speculative and further research is needed to pinpoint the underlying causes.

### Sex and gender associations of odors and embodiment illusions

Gender is of course a cultural construct, and cosmetic scents with specific gender associations are culturally bound and reinforced by branding (Lindqvist, 2013; Zellner et al., 2008). Arguably, however, the different chemical constitutions of body (sweat) odors between biological sexes might be more genetically determined, though gender-related cultural habits might indeed play a role in the chemical constitution of body odors (Havlicek & Lenochova, 2008). As mentioned in the introduction, there are differences between sexes in the composition of sweat (Penn et al., 2007), axillary skin microflora (Jackman & Noble, 1983), and non-volatile odor-precursors (Troccaz et al., 2009). In fact, while there are environmental factors playing a role in body odors (Havlicek & Lenochova, 2008), it has been argued that genetics are the primary source of such determinants (Havlicek & Lenochova, 2008; Porter, 1998; Porter et al., 1985). However, the full composition of human body volatiles is inconclusive due to methodological differences between studies (Dormont et al., 2013), and our capacity to discriminate sex through body odors remains elusive (Mutic et al., 2016). Still, this distinction between more genetically determined versus more culturally determined factors might underlie the differences between gender cosmetic scents and sex sweat odors in our study. To further support this distinction, it has been argued that body odors are of functional relevance in some cultures due to the linguistic diversity referring to such odors (Arshamian et al., 2020); by contrast, in Western culture people rather seek to hide these body odors (Perl et al., 2020). Indeed, our measures suggest that participants found significantly different gender associations between the cosmetic scents, but no differences were found between the sweat odors, which indeed follows culturally determined associations. Remarkably, it has been suggested that stronger and less pleasant body odors tend to be related to a male sex category (Doty et al., 1978), which may explain why both male and female axillary sweat have been explicitly associated to rather masculine scents (see Mutic et al., 2016).

Interestingly, no differences between conditions were found for the item regarding the gender of the own body in the experimental procedure (i.e., *Sometimes it felt like I had a male/female body*). A recent study showed that by embodying a different sex body, participants’ own gender identity is modulated in the direction of the gender associated to the embodied body (Tacikowski et al., 2020). However, a more recent study showed no modulation of the gender identity by a similar intervention, but, in contrast to our study, it did show relatively high responses regarding the feeling that they embodied the gender associated with the embodied avatar (Bolt et al., 2021). Notably, we did not assess how participants rated the gender-identity of their own bodies at baseline but only between conditions, preventing us from assessing whether there was any variation from the usual gender identification of their body. However, their ratings were relatively neutral overall (see Table 1), which might suggest a potential change from their general gender identity towards a more neutral one. Furthermore, we only addressed explicit gender identification but not implicit gender associations. There is evidence that olfactory cues implicitly communicate sex-related information of a seen body (Ye et al., 2019; Zhou et al., 2014), and future studies are encouraged to also assess implicit measures of gender identity associated with embodiment and the sense of smell. It should be noted, however, that our question on gender identity may have been ambiguous as it could have been interpreted it item in terms the visual features of the seen body rather than in terms of the identity of the own body.

In our study, the low explicit gender ratings might relate to the fact that both (female and male) actors were dressed in a similar, arguably neutral manner (Figure 1), which contrasts with previous studies wherein stereotypical visual cues were highlighted to represent the avatars’ gender (e.g., wearing a dress for female avatars while jeans and a shirt for male; Bolt et al., 2021). Notably, most bodily illusions are considered to be visually-driven (Roel Lesur et al., 2018), and perhaps the effect of our stimulation procedure could be amplified by more salient stereotypical gender visual qualities such as clothing. It could be speculated that the explicitly recognized gendered cosmetics could have stronger effects when embodying visually stereotypical bodies.

### Limitations, considerations, and outlook

Our protocol included a gesture through which participants approached their own skin to their nose to accentuate the association of the smell to the seen body, however it did not seem to help participants associate the odors with the seen body nor to attend the odors. It should be noted that participants did not recall feeling that the smells were emanating from their own bodies, despite our inclusion of the armpit sniffing part, but rather from the experimenter or the environment. However, this question was only asked after the embodiment procedure. In fact, participants recollection of having smelled the odors during the procedure was not above chance except for the female cosmetic scent. A vast amount of literature suggests that explicit recollection of odors is not as good as are implicit behaviors suggesting recollection (Degel & Köster, 1999; Herz & Engen, 1996; Zucco, 2003). As a result, such responses might be related to poor memory retrieval. In any case, it remains unclear whether the perceived source of the odors would make a strong difference, particularly given the arguably that we only found differences in embodiment at an implicit level. Future studies might want to make sure that the presented odors appear to emanate from the own (or seen) body, e.g., by repeating such body movements to the nose and only present the odor time-locked to the movement using an olfactometer rather than continuously as done in our study. In line with this, the temporal synchrony of multisensory cues has shown to be fundamental for embodiment (Botvinick & Cohen, 1998; Roel Lesur, Weijs, et al., 2020; Shimada et al., 2014). Additionally, as mentioned above, potentially enhancing stereotypical gender cues might yield different results at an explicit level.

We found differences using reconstructed sweat odors (only three compounds), based on the results of Troccaz et al. (2009). However, this is only a first approach because the content of sweat is much richer (Starkenmann, 2017) and it is likely that other compounds are involved in sexual dimorphism of sweat odors. Using a more ecological approach, it might be interesting to address this issue using real sweat samples collected from men and women to confirm our results. However, it remains important to disentangle the basic components influencing sex-related processes to elucidate the biological mechanisms our results.

Finally, it is worth noting that the way in which we studied stereotypical gender- and sex-congruencies (i.e., using certain cosmetics typically associated with the male/female sex) does not necessarily nor truthfully reflect the richness of gender expression and its dissociation from sex. Our reliance on such typical associations was for the purpose of experimental simplicity, and we by no means want to perpetuate these as normative associations.

### Conclusions

Our study highlights the relevance of olfaction in the study of bodily self-consciousness. Despite the relative neglect of this sensory modality in the field, we here show its importance for modulating implicit aspects of embodiment. This implicit modulation follows unique characteristics of the sense of smell in human cognition. Future research is needed to understand the dynamics underlying our findings more thoroughly.

## Acknowledgements

We want to thank Jasmin Schilling for her support with data collection and management, and Irene Madoka Scheiwiller for her help with piloting and preparation of the experiment. Anna Khazina, Kelly Gibbs and François-Rémy Ringeval and Tonino Rizzo for their participation in the immersive videos. The authors also thank all the members of the Human Perception and Bioresponses Department of the Research and Development Division of Firmenich, S. A., for their invaluable advice and technical competence. This project was funded by the University of Geneva–University of Zurich Joint Seed Money Funding 2018. M. R. L. and B. L. were funded by the Swiss National Science Foundation (PP00P1_170511), and Y. S. was supported by an Early Postdoc. Mobility fellowship from the Swiss National Science Foundation (P2GEP1_187911).

