## Supplementary materials for "Different armpits under my new nose: olfactory sex but not gender affects implicit measures of embodiment"

Table S1.

*Questionnaire applied for each olfactory stimulus during the Odor perception block*

| Question | Measure | Category |
|---|---|---|
| Was the presented scent female or male? | VAS, ranging from *female* (0) to *male* (1) | Gender attribution |
| Was the presented odor female or male? | Forced choice: *female/male* | Gender attribution (forced) |
| I am certain about my previous choice | Forced choice: *yes/no* | Gender attribution certainty |
| This scent was presented previously to me during the experiment | Forced choice: *yes/no* | Recognition from experiment |
| This is a typical sweat odor | Forced choice: *yes/no* | Typicality |
| I like this smell | VAS, ranging from *strongly disagree* (0) to *strongly agree* (1) | Liking |
| This scent is intense | VAS, ranging from *strongly disagree* (0) to *strongly agree* (1) | Intensity |
| I am familiar with this smell | VAS, ranging from *strongly disagree* (0) to *strongly agree* (1) | Familiarity |

Table S2

*Descriptive statistics of the embodiment questionnaire (N = 22).*

| | Congruent sweat odor | | Congruent cosmetic scent | | Incongruent sweat odor | | Incongruent cosmetic scent | |
|---|---|---|---|---|---|---|---|---|
| Item | Median | IQR | Median | IQR | Median | IQR | Median | IQR |
| I felt comfortable in the room. | 0.74 | 0.16 | 0.81 | 0.27 | 0.72 | 0.32 | 0.81 | 0.21 |
| Sometimes the body I saw when looking down felt like a stranger | 0.81 | 0.31 | 0.80 | 0.35 | 0.83 | 0.23 | 0.78 | 0.25 |
| Sometimes it felt like I could control the seen body like my own. | 0.61 | 0.42 | 0.66 | 0.45 | 0.59 | 0.35 | 0.62 | 0.46 |
| Sometimes it felt as if the touches I felt were caused by the ones I saw | 0.60 | 0.44 | 0.55 | 0.29 | 0.58 | 0.37 | 0.51 | 0.52 |
| Sometimes it felt as if the body I saw was like my own in terms of shape or skin color or appearance | 0.31 | 0.43 | 0.30 | 0.52 | 0.37 | 0.51 | 0.34 | 0.49 |
| Sometimes it felt like I was in the middle of the action. | 0.25 | 0.53 | 0.18 | 0.50 | 0.21 | 0.46 | 0.42 | 0.54 |
| Sometimes I had the feeling that I could be threatened by what I saw | 0.60 | 0.46 | 0.52 | 0.65 | 0.49 | 0.57 | 0.55 | 0.38 |
| Sometimes it felt like I had a male/female body | 0.54 | 0.55 | 0.52 | 0.42 | 0.59 | 0.42 | 0.69 | 0.33 |
| Sometimes I felt as if I could move the seen body | 0.52 | 0.36 | 0.60 | 0.51 | 0.65 | 0.39 | 0.61 | 0.36 |
| I felt good in my body | 0.70 | 0.31 | 0.70 | 0.29 | 0.72 | 0.27 | 0.70 | 0.32 |
| Sometimes it felt like I was wearing different clothes than before the experiment | 0.71 | 0.33 | 0.70 | 0.24 | 0.65 | 0.29 | 0.66 | 0.34 |
| Sometimes the body I saw felt like my own | 0.44 | 0.38 | 0.57 | 0.33 | 0.58 | 0.38 | 0.57 | 0.42 |
| In general, the touches I saw and felt were synchronous / simultaneous | 0.80 | 0.18 | 0.78 | 0.19 | 0.80 | 0.22 | 0.76 | 0.23 |

Table S3

*Results from the linear mixed model on the skin conductance response amplitude to threat.*

| | *b* | SE | 95% CI | *t* | df | *p* |
|---|---|---|---|---|---|---|
| **Fixed effects** | | | | | | |
| Intercept | .44 | .03 | [.37, .51] | 13.26 | 22.44 | < .001 |
| Congruency | .08 | .03 | [.005, .15] | 2.46 | 13.69 | .028 |
| Odor Type | .02 | .03 | [-.04, .08] | 0.91 | 9.83 | .387 |
| Interaction | .05 | .02 | [.01, .09] | 2.82 | 234.77 | .005 |
| **Random effects** | | | | | | |
| Participants | | | | | | |
| Intercept | | | 0.12 | | | |
| Congruency | | | 0.10 | | | |
| Odor Type | | | 0.08 | | | |
| r(Intercept, Congruency) | | | .13 | | | |
| r(Intercept, Odor Type) | | | -.41 | | | |
| r(Congruency, Odor Type) | | | -.45 | | | |
| Residuals | | | 0.25 | | | |

*Note.* The values reported for the random effects correspond to the standard deviation of the random effect in question or the correlation between random effects, respectively. r = correlation.

### Selection of non-bodily odors

Five members of our laboratory naive to this study rated six synthesized scents from 1 to 6, where 1 is the least pleasant and 6 the most pleasant odor. The scents were respectively rose (M = 5.2, SD = 0.75), lily (M = 5.2, SD = 0.75), soap (M = 5.2, SD = 0.75), ocean (M = 5.2, SD = 0.75), grass (M = 5.2, SD = 0.75), and mushroom (M = 5.2, SD = 0.75). The highest and the lowest scores were selected for the positive and negative stimulus respectively.
